# To β or not to β: Lack of correlation between APC mutation and β-catenin nuclear localization in colorectal cancer

**DOI:** 10.1101/2022.05.24.493182

**Authors:** Pratyusha Bala, Padmavathi Kavadipula, Sanjana Sarkar, Murali Dharan Bashyam

**Author notes:** Corresponding author: Dr Murali Dharan Bashyam, Laboratory of Molecular Oncology, Centre for DNA Fingerprinting and Diagnostics, Hyderabad 500039. Department of Medical Oncology, Dana-Farber Cancer Institute, Boston, Massachusetts Institute of Technology and Harvard University, Cambridge, MA, USA. Equal contribution (Padmavathi Kavadipula, Sanjana Sarkar).

## Abstract

Colorectal cancer (CRC) appears to arise from sequential genetic lesions in tumor suppressor genes (*APC, SMAD4* and *TP53*) and oncogenes (*KRAS*) leading to the classical adenoma to carcinoma progression. Biallelic *APC* inactivating genetic aberrations are detected in about 70% of early microadenomas implicating APC inactivation as the first genetic hit in CRC. APC is an essential protein of the Wnt ‘destruction complex’; APC inactivation is believed to cause disruption of the complex allowing stabilization and nuclear translocation of β-catenin, resulting in transcriptional activation of cancer-promoting genes. Here, we provide evidence for a surprising lack of correlation between *APC* mutation and β-catenin nuclear localization in early-onset sporadic rectal cancer samples. β-catenin nuclear localization and *APC* mutation were validated from serial FFPE sections representing same tumor regions. Several factors including status of *KRAS* mutation could not explain the absence of β-catenin nuclear localization in tumor samples harboring *APC* mutation. This lack of correlation was validated in CRC cell lines harboring various APC mutations. Our results provide evidence directly from tumor samples for possible non-canonical role(s) for mutant *APC*.

## Introduction

The tightly regulated Wnt signaling pathway co-ordinates adult stem cell division to replace epithelial cells that are continuously lost from the intestinal lumen. The cascade is activated upon binding of Wnt ligand to its cognate receptor(s) causing destabilization of a destruction complex whose primary role is to target cytoplasmic β-catenin for degradation. The stabilized β-catenin translocates into the nucleus to co-operate with TCF/LEF transcription factor family members causing activation of pro-colorectal cancer (CRC) genes (1). Adenomatous Polyposis Coli (APC), encoded by the *APC* gene, constitutes an important scaffold of the destruction complex and its mutational inactivation forms the basis of the central dogma of CRC. APC interacts with and regulates β-catenin via specific domains including three 15 amino-acid repeats (15AARs, between codons 1021 and 1170), seven 20 amino-acid repeats (20AARs, between codons 1265 and 2035) and seven armadillo repeats (ARM, codons 453– 767) (2). Of these, the seven 20AARs are believed to be more important and are cardinal with respect to *APC* mutation in CRC (3). Majority of inactivating mutations occur in a ‘mutation cluster region’ (MCR) of APC defined by amino acid positions 1286-1514 (4) and lead to a truncated protein that loses most (but not all) of the seven 20AARs (5). It has been suggested that loss of all seven repeats is detrimental to cell survival (6). However, the position of APC truncations and the consequent effect on β-catenin status has been more robustly studied for the familial form of CRC called familial adenomatous polyposis (FAP), with an etiology similar to Wnt-dependent sporadic CRC (7). In FAP, there is a mono-allelic germline inactivation of *APC* and tumors develop following a somatic inactivation of the remaining allele (8).

APC inactivation is believed to result in constitutive activation of β-catenin transcription function in the absence of stimuli from the Wnt ligand. β-catenin (or plakoglobin) is known to anchor E-cadherin on the cytoplasmic side of the membrane to the actin cytoskeleton through α-catenin (9). Immunohistochemistry (IHC)-based evaluation of β-catenin nuclear vs cytoplasmic/membrane localization is considered the hallmark for evaluation of Wnt status in CRC; nuclear localization indicating activation of Wnt signaling (10). Research spread over past four decades has validated the high frequency (≥ 70%) of *APC* mutations in sporadic CRC (4,11). However, concomitant identification of β-catenin nuclear localization status is often lacking in these studies.

## Materials and methods

### Samples

The study was performed following approval of ethics committee of hospitals from where samples were collected as well as of the CDFD bioethics committee as per the revised Helsinki declaration. Sample collection (following informed consent), storage, processing and isolation of DNA are described in our previous studies (12). Exome sequencing was performed as described earlier (13).

### β-catenin immunohistochemistry (IHC)

IHC was utilized to determine localization status (nuclear or membrane/cytoplasmic) of β-catenin as described earlier (12). Briefly, 4-μm sections from formalin fixed paraffin embedded (FFPE) blocks were de-paraffinized in Xylene and hydrated in a graded series of alcohol solutions. Heat-induced antigen retrieval was performed by pressure cooker retrieval method in citrate buffer (1 mM, pH 6) before staining. Endogenous peroxidase was blocked by pre-incubation with 2 % Hydrogen Peroxide in methanol. The sections were then incubated with anti β-catenin antibody (BD Biosciences, California, United States) for 1 hour at room temperature followed by treatment with HRP-conjugated anti-mouse secondary antibody (DAKO Envision Plus HRP kit, Dako Denmark A/S, Glostrup, Denmark) for 30 minutes. Sections were then subjected to chromogenic detection using DAB reagent from the Envision Plus HRP kit and were counter-stained with haematoxylin. The β-catenin status was scored as positive if the stain was observed in the nuclei of more than 30% of tumor epithelial cells and negative if the stain was detected in less than 20% of cells; samples with intermediate stain (20-30% nuclear stain) were excluded from the analysis.

### DNA isolation from FFPE

DNA was isolated from FFPE samples for APC mutation validation. 8 μm FFPE sections were cut using a microtome (Leica RM2125 RT, Leica Biosystems, Germany) and fixed onto frosted micro slides (Blue Star, Mumbai, India). Tumor-rich areas were marked by the pathologist on a haematoxylin-stained section and tumor cells were manually micro-dissected from the marked regions under a microscope. DNA was isolated from the micro-dissected tumor cells using SDS/proteinase K digestion at 58°C overnight, followed by standard phenol-chloroform extraction and alcohol precipitation. Isolated DNA was quantitated using a spectrophotometer and stored at -20°C.

### Cell culture

Human CRC cell lines HCT116, HT29, COLO205, Caco-2, DLD1, HCT15, SW620 and SW480 (procured from National Center for Cell Science, Pune, India), authenticated via STR profiling and confirmed to be free of mycoplasma contamination, were maintained in Dulbecco’s Modified Eagle’s Medium (DMEM) supplemented with 10% FBS and antibiotic/antimycotic solution (ThermoFisher Scientific, Waltham, MA, USA).

Cells were grown on chamber slides (BD Biosciences) for 24hr, fixed in 4% paraformaldehyde for 15min at room temperature followed by permeabilization using 0.1% Triton X 100 for 10min. Cells were washed with PBS and blocking was done in 5% (w/v) bovine serum albumin for 1hr at room temperature. After blocking, the fixed material was incubated overnight with β-catenin primary antibody (BD Biosciences) at 4°C. Cells were washed with PBS for 5 min followed by incubation with Alexa fluor-488 conjugated secondary antibody (ThermoFisher Scientific) for 1hr at room temperature. Cells were then mounted using Vectashield mounting medium with DAPI and visualized with LSM 700 confocal laser scanning microscope (Carl Zeiss, Oberkochen, Germany) or confocal microscope (Leica SP-8, Wetzlar, Germany) at 63X magnification.

## Results

Our previous studies revealed the absence of aberrant Wnt/β-catenin signaling in a significant proportion of early onset sporadic rectal cancer (EOSRC), the predominant CRC subtype in India (12). We recently reported identification of *APC* mutation, via exome sequencing analysis, in 22 of 47 carefully selected and well annotated EOSRC samples (13). The reduced frequency of APC mutations, compared to other reports in the literature, appeared to validate our previous observations pertaining to reduced frequency of aberrant Wnt activation in EOSRC. However, in order to establish a causal relationship between the presence of *APC* mutation and status of Wnt signaling, we determined the intracellular localization status of β-catenin using IHC. Surprisingly, only 10 of the 22 samples harboring a mutant APC, exhibited β-catenin nuclear localization (Table 1).

**Table 1:** Analysis of *APC* mutation and β-catenin nuclear localization status in EOSRC tumor samples.

| Sample ID | APC cDNA_change* | APC protein_change | Variant class* | Strelka_VAF* | Mutect2_VAF* | Intact 20AARs remaining in mutant APC | β-catenin nuclear localization status | KRAS mutation status* |
| --- | --- | --- | --- | --- | --- | --- | --- | --- |
| 1095 | c.3632T>G | p.Met1211Arg | MS | 0.57 | 0.58 | 7 | P | MUT |
| 2669 | c.3871C>T;<br>c.7645C>T | p.Gln1291*;<br>p.Arg2549Cys | MS | 0.38;<br>0.7 | 0.36;<br>0.65 | 7 | P | WT |
| 2215 | c.4260delC | p.Ser1421fs | FS | 0.5 | 0.44 | 2 | P | MUT |
| 2643 | c.4327_4402dup76 | p.Lys1468fs | FS & NS |  | 0.5 | 2 | P | WT |
| 2769 | c.4469delA | p.His1490fs | FS | 0.28 | 0.2 | 2 | P | MUT |
| 2827 | c.2626C>T;<br>c.4473dupT | p.Arg876*;<br>p.Ala1492fs | FS | 0.32;<br>0.46 | 0.33;<br>0.41 | 2 | P | MUT |
| 2265 | c.3916G>T | p.Glu1306* | NS | 0.25 | 0.26 | 1 | P | WT |
| 1097 | c.3766C>T | p.Gln1256* | NS | 0.3 | 0.27 | 0 | P | WT |
| 2911 | c.637C>T | p.Arg213* | NS | 0.53 | 0.51 | 0 | P | WT |
| 2299# | c.1600_1609delAAATCTGAAA;<br>c.1614_1622delAGACTTACA | p.Lys534fs;<br>p.Glu538_Gln542del | FS;<br>SIV | 0.5;<br>0.5 | 0.53;<br>0.57 | 0 | P | WT |
| 2389 | c.4682dupA | p.Asp1562fs | FS | 0.51 | 0.53 | 3 | N | WT |
| 2397 | c.4729G>T | p.Glu1577* | NS | 0.45 | 0.45 | 3 | N | MUT |
| 2455 | c.4179dupT | p.Asp1394fs | FS & NS | 0.18 | 0.21 | 2 | N | MUT |
| 2303 | c.4057G>T | p.Glu1353* | NS | 0.37 | 0.33 | 1 | N | MUT |
| 2513 | c.3924dupA | p.Glu1309fs | FS | 0.46 | 0.43 | 1 | N | WT |
| 2577 | c.3864delA;<br>c.4738_4741delATTT | p.Cys1289fs;<br>p.Ile1580fs | FS;<br>FS | 0.34;<br>0.26 | 0.27;<br>0.29 | 1 | N | MUT |
| 2667 | c.4012C>T | p.Gln1338* | NS | 0.43 | 0.41 | 1 | N | MUT |
| 2699 | c.3871C>T | p.Gln1291* | NS | 0.51 | 0.49 | 1 | N | WT |
| 2729 | c.4037C>A | p.Ser1346* | NS | 0.45 | 0.46 | 1 | N | MUT |
| 2737 | c.1495C>T | p.Arg499* | NS | 0.21 | 0.22 | 0 | N | WT |
| 2775 | c.637C>T | p.Arg213* | NS | 0.46 | 0.45 | 0 | N | WT |
| 2785 | c.2804dupA | p.Tyr935fs | FS & NS | 0.76 | 0.77 | 0 | N | WT |

VAF, variant allele frequency; MS, missense variant; FS, frameshift variant; NS, stop gained variant; SIV, structural interaction variant; N, nuclear negative; P, nuclear positive; AARs, amino acid repeats.
#, 2299 sample has 2 monoallelic APC mutations.
\*, from ref 13

One possible explanation for the discrepancy between *APC* mutation and β-catenin nuclear localization could be that the cells harboring *APC* mutation represented a very small fraction of tumor epithelium and therefore may not contribute significantly to β-catenin status of the bulk tumor. However, the mutant *APC* read fraction was substantially high for all 22 samples with no significant difference between samples that did or did not exhibit β-catenin nuclear localization (Table 1). Another possible explanation for this surprising result could be that the *APC* mutation and immunohistochemistry represented different regions of the same tumor sample (intra-tumor heterogeneity). We therefore evaluated *APC* mutation and β-catenin localization status using serial sections from the same FFPE block. As shown in Figure 1, we were able to validate the *APC* mutation and β-catenin status in sections obtained from the same FFPE block and representing the same area of the tumor sample.

**Figure 1:**
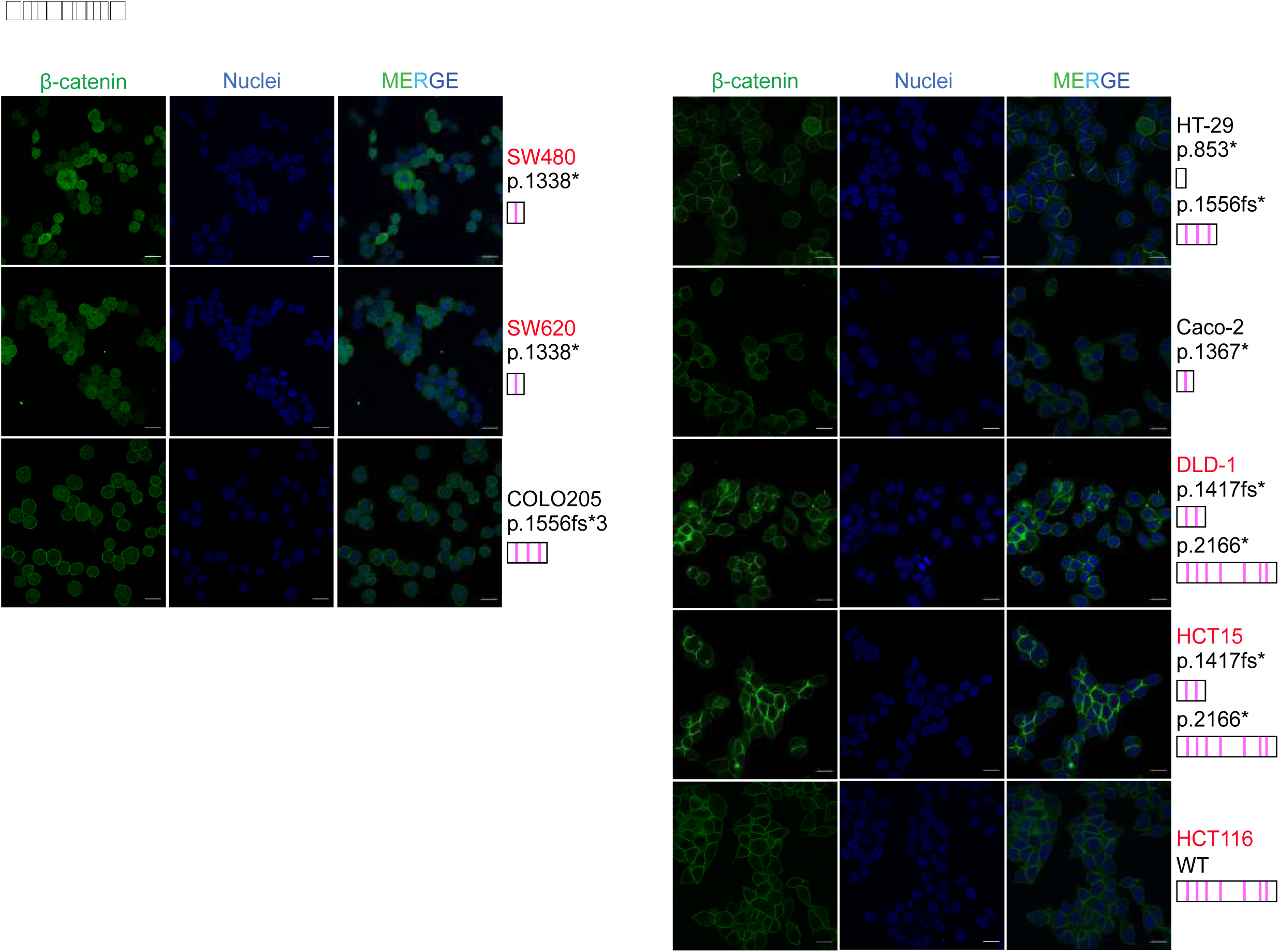
Determination of β-catenin nuclear localization status in tumors harboring *APC* mutations (indicated). Immunohistochemistry images and sequence electropherograms are shown for four *APC* mutant EOSRC tumor samples, three not showing and one exhibiting nuclear localization of β-catenin. Scale bars are 100μM.

*APC* mutations are under strict selection in CRC; the complete absence of *APC* is thought to be detrimental to a healthy colonic epithelium (6) and is expected to be unsuitable for CRC progression as well. According to the ‘just right’ signaling model (14), the truncated *APC* usually retains a few of the seven 20AARs. More importantly, the number of 20AARs that are remaining due to the *APC* truncation mutation may determine the ‘quantum’ of β-catenin nuclear localization. Therefore, we explored the possibility that absence of β-catenin nuclear localization in EOSRC tumor samples despite the presence of mutant APC, was due to the presence of sufficient β-catenin binding domains in the truncated APC. As per previous reports (15) one would expect ≥ three of the seven 20AARs to be present in samples not exhibiting β-catenin nuclear positivity. However, as shown in Table 1, there was no significant difference in the number of 20AARs remaining in the truncated APC, between β-catenin nuclear positive and negative samples.

Another factor expected to have a bearing on β-catenin nuclear localization is the status of the remaining *APC* allele. Interestingly, except three that exhibited bi-allelic APC mutations, the rest of the 19 samples harbored a wild type second allele (based on exome and sanger sequencing analysis), also indicating the absence of loss of heterozygosity. Unfortunately, we could not determine the status of DNA methylation induced possible inactivation of the second allele in samples exhibiting a mono-allelic APC mutation. More importantly, even among the three samples exhibiting bi-allelic APC mutation, there was no correlation between β-catenin nuclear localization status and the position of APC truncation (Table 1).

CRC cell lines are frequently used to study Wnt signaling. We therefore proceeded to determine the intracellular localization of β-catenin in eight carefully chosen CRC cell lines (Figure S1), of which seven exhibited various mutant forms of APC (while the eighth (HCT116) exhibited wild type APC). Surprisingly, only three of the seven exhibited β-catenin nuclear localization (Figure S1); similar to our observations from tumor samples. Of note, three cell lines (DLD1, HCT15 and HT29) harbored bi-allelic APC mutations, but none of these exhibited β-catenin nuclear localization.

Mutational activation of *KRAS* is regarded as the ‘second hit’ that follows *APC* mutation during the classical ‘CRC adenoma to carcinoma progression’ (16). It is also thought that a gain of function *KRAS* mutation may be required to facilitate β-catenin nuclear localization in colorectal tumors harboring truncated APC (17). We therefore analyzed *KRAS* mutation frequency in the 22 *APC* mutant samples; 4/10 (40%) exhibiting β-catenin nuclear localization harbored *KRAS* mutation while 6/12 (50%) samples devoid of β-catenin nuclear localization harbored *KRAS* mutation (Table 1). Similarly, 2/4 *APC* mutant cell lines devoid of β-catenin nuclear positivity, exhibited *KRAS* mutation (Figure S1). Thus, *KRAS* mutation status could not explain absence of β-catenin nuclear localization in colorectal cancer tumors or cell lines harboring *APC* mutation.

## Discussion

*APC* mutations leading to the activation of β-catenin oncogenic functions constitutes the central dogma of CRC. Here, we show that *APC* mutations may not correlate with the nuclear translocation of β-catenin in CRC. Of interest, several earlier studies revealed the absence of β-catenin nuclear translocation despite bi-allelic *APC* mutation in FAP (18,19). Moreover, recent studies reveal that mutated/truncated APC may still associate with the destruction complex even with loss of all AXIN2 (an important scaffold protein of the destruction complex) and β-catenin binding sites (20,21). A more recent study (22) showed that truncated APC mutants T1556* and S811* (that harbored none of the seven beta catenin binding domains) could still participate in the formation of an active destruction complex and successfully recruited β-catenin suggesting that the APC 20AARs might not be an exclusive supporter of β-catenin degradation. Are other APC domains also crucial in determining fate of β-catenin (23)? Our own results reveal the absence of β-catenin nuclear localization despite *APC* mutation in CRC tumors and cell lines suggesting an alternative mode of inhibition of β-catenin nuclear localization by truncated APC. Moreover, even an oncogenic *KRAS* mutation in the background of truncated APC, did not appear to trigger β-catenin nuclear localization. Thus, nuclear translocation of β-catenin may depend on factors additional to truncated APC.

However, if an *APC* mutation is under positive selection in the tumor but does not result in β-catenin nuclear localization; does it perform additional (non-canonical) roles in driving CRC? Several previous studies have suggested role(s) for APC unrelated to β-catenin regulation (reviewed in 22) including the latest discovery of its possible function in post-translational modifications based on a Drosophila proteomic screen (24). Earlier studies have suggested APC’s ability to regulate mitotic spindle contraction (25).

In a nutshell, we provide perhaps the first evidence from patient tumor samples, of the absence of a correlation between APC truncation and β-catenin nuclear translocation, pointing towards possible non-canoncial APC functions in CRC. Thus, the role of *APC* mutations in CRC does not appear to be as straightforward as was thought to be, at least in sporadic cases. It was more than three decades ago when APC was identified as a tumor suppressor in CRC and aberrant Wnt/β-catenin activation was proposed as the driver of oncogenesis. However, despite several clinical trials, no Wnt-based therapy is approved, indicating perhaps the importance of non-canonical roles of APC in CRC (Table S1). Further work needs to be performed to identify the exact role(s) of mutant *APC* in CRC.

## Author Contributions

MDB and PB conceived the study; PB and PK performed the experiments; MDB, SS, PB and PK analysed and compiled the data; MDB, PB and SS wrote the manuscript.

## Acknowledgements

We thank all patients for kindly agreeing to be a part of the study. We are grateful to Mr Viswakalyan Kotapalli, CDFD, Hyderabad, India, for advice on immunohistochemistry and imaging of stained tumor sections. We are grateful to Dr Swarnalata Gowrishankar, Apollo Hospitals, Hyderabad, India and Dr Satish Rao, Krishna Institute of Medical Sciences, Hyderabad, India, for evaluation of β-catenin immunohistochemistry slides. We thank Sara A George for correcting the manuscript. We acknowledge the Sophisticated Equipment Facility, CDFD, Hyderabad, India, for fluorescence microscopy and Sanger sequencing. The work was supported by a grant (SB/SO/HS-007/2013) from the Department of Science and Technology, Government of India to MDB.

## Figure legends

**Supplementary Figure 1:**
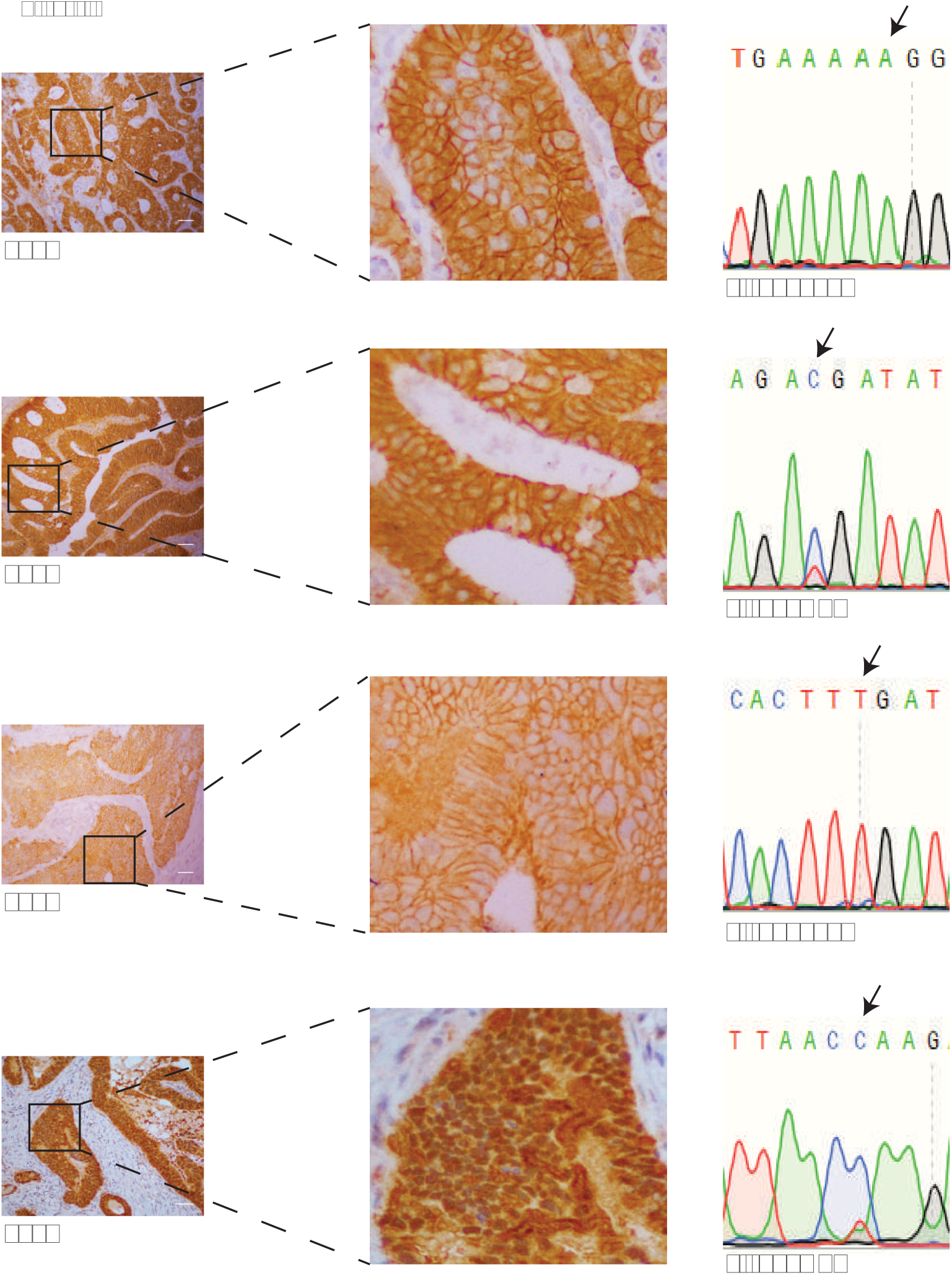
Evaluation of β-catenin localization in CRC cell lines with and without APC mutation (indicated). Status of the APC 20 AARs are indicated by vertical pink colour bars separately for each mutation (HCT116 harbors a wild type APC and so all seven AARs are intact). Cell line labels are colour coded to indicate *KRAS* status (mutant in red and wild type in black). Scale bars are 20μM.

---

**Table S1:** Clinical trials for Wnt pathway inhibitors.

| Drug | Mechanism of action | Phase | Identifier* |
| --- | --- | --- | --- |
| WNT974 (with LGX818 and Cetuximab) | PORCN inhibitor | Phase 1 | NCT02278133 |
| WNT974 | PORCN inhibitor | Phase 2 | NCT02649530 |
| ETC-1922159 | PORCN inhibitor | Phase 1 | NCT02521844 |
| RXC004 | PORCN inhibitor | Phase 1 | NCT03447470 |
| CGX1321 | PORCN inhibitor | Phase 1 | NCT03507998 |
| CGX1321 (with Pembrolizumab) | PORCN inhibitor | Phase 1 | NCT02675946 |
| OTSA101-DTPA-90Y | FZD10 antagonist | Phase 1 | NCT01469975 |
| OMP-18R5 | Monoclonal antibody against FZD receptors | Phase 1 | NCT01345201 |
| OMP-54F28 | FZD8 decoy receptor | Phase 1 | NCT01608867 |
| PRI-724 | CBP/β-catenin antagonist | Phase 1 | NCT01302405 |

PORCN, Protein-serine O-palmitoleoyltransferase porcupine; FZD, Frizzled protein; CBP, CREB binding protein
\*, https://clinicaltrials.gov/

## References

1. Aberle H, Bauer A, Stappert J, Kispert A, Kemler R. Catenin Is a Target for the Ubiquitin-Proteasome Pathway. EMBO J. 1997;16(13):3797–804.

2. Näthke IS. The Adenomatous Polyposis coli protein: The achilles heel of the gut epithelium. Annu Rev Cell Dev Biol. 2004;20:337–66.

3. Eklof Spink K, Fridman SG, Weis WI. Molecular mechanisms of beta-catenin recognition by adenomatous polyposis coli revealed by the structure of an APC-beta-catenin complex. EMBO J. 2001;20(22):6203–6212.

4. Mori Y, Nagse H, Ando H, Horii A, Ichii S, Nakatsuru S, et al. Somatic mutations of the APC gene in colorectal tumors: Mutation cluster region in the APC gene. Hum Mol Genet. 1992;1(4):229–33.

5. Kohler EM, Derungs A, Daum G, Behrens J, Schneikert J. Functional definition of the mutation cluster region of adenomatous polyposis coli in colorectal tumours. Hum Mol Genet. 2008;17(13):1978–87.

6. Browne SJ, Williams AC, Hague A, Butt AJ, Paraskeva C. Loss of apc protein expressed by human colonic epithelial cells and the appearance of a specific low-molecular-weight form is associated with apoptosis IN vitro. Int J Cancer. 1994;59(1):56–64.

7. Kinzler, Kenneth W, Vogelstein B. Lessons from Hereditary Colorectal Cancer. Cell. 1996;87:159–70.

8. Fearnhead NS, Britton MP, Bodmer WF. The ABC of APC. Hum Mol Genet. 2001;10(7):721–33.

9. Yap AS, Brieher WM, Gumbiner BM. Molecular and functional analysis of cadherin-based adherens junctions. Annu Rev Cell Dev Biol. 1997;13:119–46.

10. MacDonald BT, Tamai K, He X. Wnt/β-Catenin Signaling: Components, Mechanisms, and Diseases. Dev Cell [Internet]. 2009;17(1):9–26.

11. Palacio-Rúa KA, Isaza-Jiménez LF, Ahumada-Rodríguez E, Ceballos-García H, Muñetón-Peña CM. Genetic analysis in APC, KRAS, and TP53 in patients with stomach and colon cancer. Rev Gastroenterol Mex. 2014;79(2):79–89.

12. Raman R, Kotapalli V, Adduri R, Gowrishankar S, Bashyam L, Chaudhary A, et al. Evidence for possible non-canonical pathway(s) driven early-onset colorectal cancer in India. Mol Carcinog. 2014;53(S1):181–6.

13. Bala P, Singh AK, Kavadipula P, Kotapalli V, Sabarinathan R, Bashyam MD. Exome sequencing identifies ARID2 as a novel tumor suppressor in early-onset sporadic rectal cancer. Oncogene [Internet]. 2021;40(4):863–74.

14. Albuquerque C, Breukel C, Van Der Luijt R, Fidalgo P, Lage P, Slors FJM, et al. The “just-right” signaling model: APC somatic mutations are selected based on a specific level of activation of the β-catenin signaling cascade. Hum Mol Genet. 2002;11(13):1549–60.

15. Munemitsu S, Albert I, Souza B, Rubinfeld B, Polakis P. Regulation of intracellular beta-catenin levels by the adenomatous polyposis coli (APC) tumor-suppressor protein. Proc Natl Acad Sci U S A. 1995;92(7):3046–3050.

16. Janssen KP, Alberici P, Fsihi H, et al. APC and oncogenic KRAS are synergistic in enhancing Wnt signaling in intestinal tumor formation and progression. Gastroenterology 2006; 131:1096–1109.

17. Phelps RA, Chidester S, Dehghanizadeh S, et al. A two-step model for colon adenoma initiation and progression caused by APC loss. Cell. 2009;137(4):623–634.

18. Bläker H, Scholten M, Sutter C, Otto HF, Penzel R. Somatic Mutations in Familial Adenomatous Polyps: Nuclear Translocation of β-Catenin Requires More Than Biallelic APC Inactivation. Am J Clin Pathol. 2003;120(3):418–23.

19. Yedid N, Kalma Y, Malcov M, Amit A, Kariv R, Caspi M, et al. The effect of a germline mutation in the APC gene on β-catenin in human embryonic stem cells. BMC Cancer [Internet]. 2016;16(1):1–13.

20. Li VSW, Ng SS, Boersema PJ, Low TY, Karthaus WR, Gerlach JP, et al. Wnt Signaling through Inhibition of β-Catenin Degradation in an Intact Axin1 Complex. Cell [Internet]. 2012;149(6):1245–56.

21. Voloshanenko O, Erdmann G, Dubash TD, Augustin I, Metzig M, Moffa G, et al. Wnt secretion is required to maintain high levels of Wnt activity in colon cancer cells. Nat Commun. 2013;4(May):1–13.

22. Ranes M, Zaleska M, Sakalas S, Knight R. Article Reconstitution of the destruction complex defines roles of AXIN polymers and APC in b -catenin capture, phosphorylation, and ubiquitylation ll Reconstitution of the destruction complex defines roles of AXIN polymers and APC in b -catenin capture. Mol Cell [Internet]. 2021;81(16):3246-3261.e11.

23. Kimelman D, Xu W. beta-catenin destruction complex: insights and questions from a structural perspective. Oncogene. 2006;25(57):7482–7491.

24. Blundon MA, Schlesinger DR, Parthasarathy A, Smith SL, Kolev HM, Vinson DA, et al. Proteomic analysis reveals APC-dependent post-translational modifications and identifies a novel regulator of β -catenin. 2016;2629–40.

25. Sugioka K, Fielmich LE, Mizumoto K, Bowerman B, Van Den Heuvel S, Kimura A, et al. Tumor suppressor APC is an attenuator of spindle-pulling forces during C. elegans asymmetric cell division. Proc Natl Acad Sci U S A. 2018;115(5):E954–63

