## Supplementary figures and images for "To β or not to β: Lack of correlation between APC mutation and β-catenin nuclear localization in colorectal cancer"

### Supplementary Figure 1

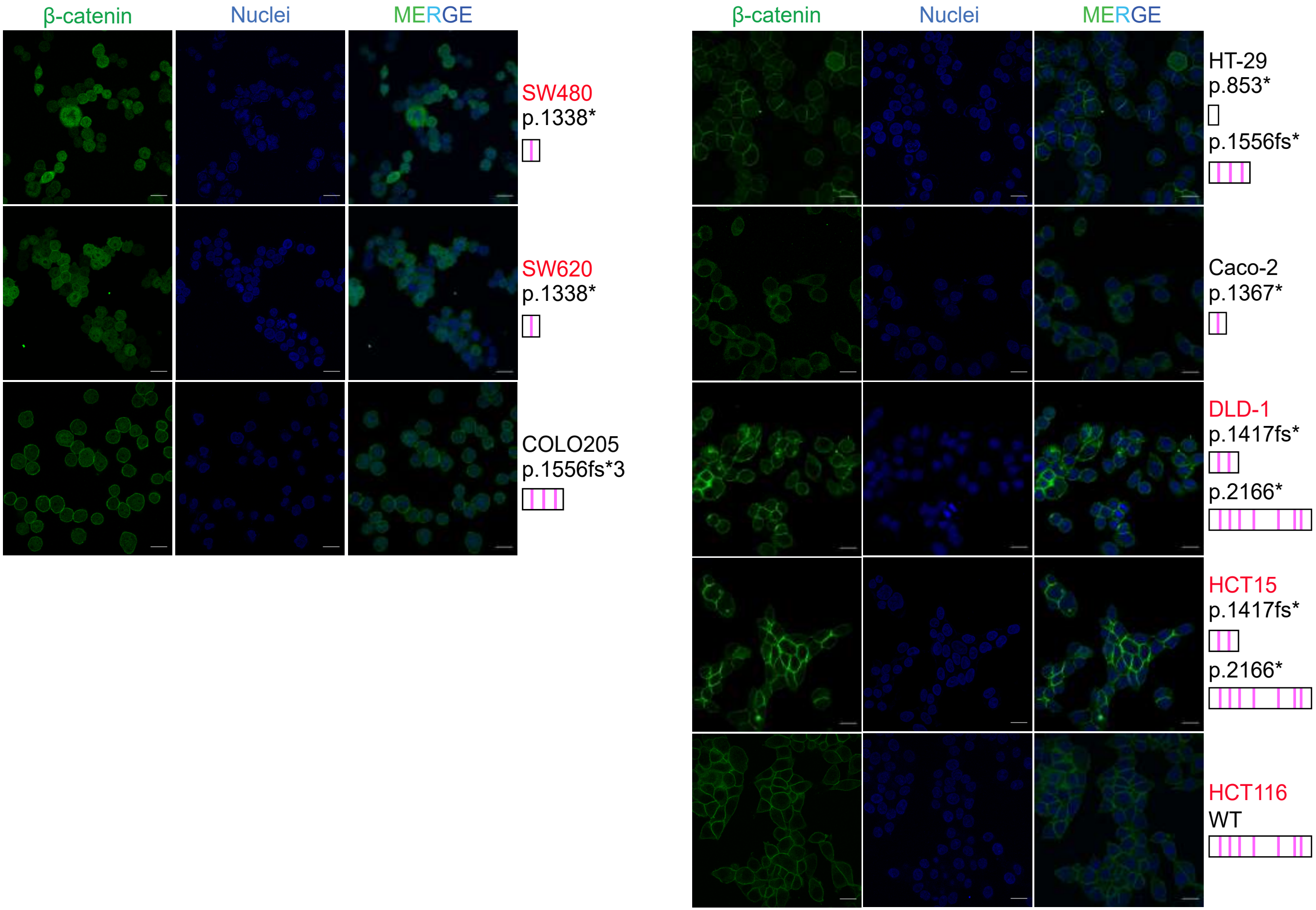
